# Deciphering the transcriptional regulatory network of *Yarrowia lipolytica* using machine learning

**DOI:** 10.1101/2024.07.29.605545

**Authors:** Abraham A.J. Kerssemakers, Jayanth Krishnan, Kevin Rychel, Daniel C. Zielinski, Bernhard O. Palsson, Suresh Sudarsan

## Abstract

The transcriptional regulatory network (TRN) in Yarrowia lipolytica coordinates its cellular processes, including the response to various stimuli. The TRN has been difficult to study due to its complex nature. In industrial-size fermenters, environments are often not homogenous, resulting in Yarrowia experiencing fluctuating conditions during a fermentation. Compared with homogenous laboratory conditions, these fluctuations result in altered cellular states and behaviors due to the action of the TRN. Here, a machine learning approach was deployed to modularize the transcriptome to enable meaningful description of its changing composition. To provide a sufficiently broad dataset, a wide range of relevant fermentation conditions (nutrient limitations, growth rates, pH values, oxygen availability and CO2 stresses) were run and samples obtained for RNA-Seq generation. We thus significantly increased the number of publicly available transcriptomic dataset on Y. lipolytica W29. In total, 23 independently modulated gene sets (termed iModulons) were identified of which 9 could be linked to corresponding regulons in S. cerevisiae. Strong responses were found in relation to oxygen limitation and elevated CO2 concentrations represented by (i) altered ribosomal protein synthesis, (ii) cell cycle disturbances, (iii) respiratory gene expression, and (iv) redox homeostasis. These results provide a fine-grained systems-level understanding of the Y. lipolytica TRN in response to industrially meaningful stresses, providing engineering targets to design more robust production strains. Moreover, this study provides a guide to perform similar work with poorly characterized single-cellular eukaryotic organisms.

**Highlights:**

- A large screening setup significantly expands the public RNA-Seq library on *Y. lipolytica*.
- This work provides a systems-level understanding of the TRN in response to industrial stresses.
- iModulon analysis can help accelerate bioprocess development.
- Results can guide similar work on poorly characterized single-cellular eukaryotic organisms.

## Introduction

*Y. lipolytica* is an oleaginous, strictly aerobic yeast that is gaining interest due to its wide substrate and product spectrum, and general robustness(Nicaud, 2012). As a result, combined with an increased genomic understanding, it is becoming a popular host for building economically viable bioprocesses(Bankar et al., 2009; Liu et al., 2015; Park and Ledesma-Amaro, 2023). However, bioprocess development brings many different challenges. One of the most dominant problems is altered strain behavior and performance when exposed to large-scale (industrial) conditions(Straathof et al., 2019). Although there are a variety of reasons for this change in behavior, it is often a stress response caused by environmental fluctuations or pressures as a cell circulates through various regimes within a bioreactor(Lara et al., 2006). Microorganisms sense these changes through varying mechanisms, and different strategies are initiated to respond to them accordingly(Mitchell et al., 2015; Seshasayee et al., 2006). In a laboratory setting, these responses are not often observed as bioreactor conditions are kept homogeneous and within an ideal range. However, to facilitate a more efficient bioprocess development, it is crucial to gain insights into complex cellular regulation. As such, scale-down experiments become very useful and informative, aiming to replicate industrial conditions in the laboratory and assess strain behavior under realistic conditions(Buchholz et al., 2014; Kerssemakers et al., 2023; Noorman and Heijnen, 2017).

Eukaryotic regulation is highly complex and the transcriptional regulatory network (TRN) plays an important role in directing cellular function(Balaji et al., 2006; Lelli et al., 2012). In response to various stimuli, the TRN defines relationships between regulators and genes(Lee et al., 2002). Although great steps have been made on understanding the model organism *S. cerevisiae*, this understanding is still limited in *Y. lipolytica*(Abdulrehman et al., 2011; Monteiro et al., 2020a). Therefore, this research deploys a systems biology approach to elucidate the TRN in *Y. lipolytica*. More specifically, it applies the concept of Independent Component Analysis (ICA) to obtain independently modulated and co regulated gene sets (iModulons) (Hyvärinen and Oja, 2000; Sastry et al., 2019). Given a sufficiently large dataset, ICA can decompose multivariate signals into independent signals and their relative strengths. Biologically speaking, this enables iModulons to capture genes regulated by the same TF, thereby gaining new insights into the TRN structure. Unlike traditional approaches, which require significant cost, effort, and time to analyze each regulator one at a time from the bottom up, ICA quantifies the effects of all sufficiently variable regulators at once, from the top-down. This approach has been applied successfully for a variety of prokaryotes in a well-developed database, iModulonDB.org(Rychel et al., 2021), with *E. coli* as primary example(Lamoureux et al., 2022).

The usefulness of ICA is directly linked to the quality and quantity of available datasets. As *Y. lipolytica* is not yet regarded as a model organism, there is limited omics analysis available. Therefore, to increase the data availability, a significant set of fermentations with subsequent RNA sequencing has been performed. In this work, the exposure to a series of relevant industrial stressors (nutrient limitations, pH, carbon sources, oxygen availability, and elevated CO_2_ levels) gives insights into the cellular rearrangements that could potentially lead to reduced performance. Outcomes can be used as a monitoring tool to assess the presence of reactor heterogeneities or aid in the development of strains with an increased robustness against the suboptimal conditions.

With this diverse and high-quality dataset, ICA was performed and yielded a set of iModulons. These results provide new insights into the complex transcriptional regulation of *Y. lipolytica*. The novel insights can be used as a design tool to facilitate more efficient bioprocess development from the laboratory to an industrial scale. As this work is the first of its kind to apply iModulon analysis on a eukaryotic strain, it also positions itself as a promising proof of concept.

## Material and methods

### Strain and media

The wild-type *Y. lipolytica* W29 and succinic acid overproducer ST8578 were used in this research(Babaei et al., 2019).

Pre-cultures were prepared by using 250 µL stock culture (25% glycerol, -80 °C) to inoculate 25 mL Delft media (20 g/L glucose, 14.4 g/L KH_2_PO_4_, 0.5 g/L MgSO_4_, 7.5 g/L (NH_4_)_2_SO_4_, 0.1 mg/L biotin, 0.4 mg/L 4-aminobenzoic acid, 2 mg/L nicotinic acid, 2 mg/L calcium pantothenate, 2 mg/L pyridoxine, 2 mg/L thiamine HCl, 50mg/L myo-inositol, 4.5 mg/L CaCl_2_·2H_2_O, 4.5 mg/L ZnSO_4_·7H_2_O, 3 mg/L FeSO_4_·7H_2_O, 1mg/L boric acid, 1 mg/L MnCl_2_·4H_2_O, 0.4 mg/L Na_2_MoO_4_·2H_2_O, 0.3 mg/L CoCl_2_·6H_2_O, 0.1 mg/L CuSO_4_·5H_2_O, 0.1 mg/L KI and 15 mg/L EDTA) and left to grow overnight at 30 °C and 225 RPM.

### Shaken cultivations in micro titer plates

A set of 24 different conditions were designed for the shaken cultures. Variations included are carbon source (glucose and glycerol), carbon loadings (20 g/L and 80 g/L), nutrient limitations (N, P and Mg), pH values (buffered and unbuffered) and oxygen availability based on the filling volumes of the deep wells. Standard Delft media was taken as reference condition. In the instance where glycerol was used, other media components remained the same. For the higher carbon loading, all components were scaled accordingly. Nutrient limitations for N, P and Mg were achieved by decreasing (NH_4_)_2_SO_4_ five-fold, KH_2_PO_4_ 28-fold and MgSO_4_ 17-fold respectively. All media compositions were brought to pH 6 and buffered cultures were supplemented with 0.1 M 2-(N-morpholino)ethanesulfonic acid (MES) buffer. Lastly, oxygen availability was adjusted by filling the 12 mL wells with either 1.5, 2.5 or 3.5 mL (“Webpage:,” n.d.).

For the pre-culture, 25 mL of Delft media was inoculated with 250 µL of glycerol stock (25% v/v at −80 °C) and grown overnight at 30 °C and 225 rpm. W29 pre-culture was used to inoculate all respective culture conditions at an OD_600_ of 0.1. Each condition was prepared in 12 mL deep well plates (Enzyscreen, the Netherlands) in duplicates. Cultivation was done in a Growth profiler (Enzyscreen, the Netherlands) at 30 °C and a shaking speed of 250 RPM. All cultures were sampled as the reference culture (1.5 mL, buffered, 20 g/L glucose, without limitations) reached mid-log phase.

### Continuous bioreactor cultivations

Continuous cultivations were performed in a 1L Biostat Q Plus vessel (Sartorius, France) with a working volume of 0.5 L. Dissolved oxygen (DO) was measured with the VisiFerm DO 160 probe (Hamilton Company, US) and controlled by adjusting the flowrate of air, and by mixing with pure O_2_ or N_2_. A minimum flow rate of 300 mL/min was used to ensure that sufficient airflow was supplied to the off-gas analyzer. Stirring was done with a single Rushton impeller at a constant speed of 500 RPM to avoid increased liquid levels due to increased agitation. pH was monitored with the EasyFerm Plus PHI k8 160 probe (Hamilton Company, US) and maintained at pH 6 by automatic addition of 5M NaOH using the built in PID control loop. The temperature was set at 30 °C and was controlled automatically through PID. Feed rates were controlled by external pumps that were calibrated prior to use. Media was designed based on the Delft composition with a glucose concentration of 9 g/L and the addition of 1 mL/L of Antifoam 204 (Merck, Germany). The bioreactors were inoculated at an OD_600_ of 0.1. After completion of the batch-phase, feed and harvest pumps were activated. Five residence times passed before reaching the first steady state, followed by two residence times for the remaining conditions.

The first continuous fermentation aimed to collect data on the profiles across different growth rates. Two cultures (strains W29 and ST8578) were operated at subsequent growth rates of 0.05, 0.1, 0.15, 0.2, 0.25, 0.3 and 0.05, 0.075, 0.1, 0.125, 0.15 respectively.

The second continuous fermentation aimed to increase dissolved CO_2_ levels through increasing the CO_2_ partial pressure in the sparging air. To achieve this, pure CO_2_ and oxygen were mixed with their individual mass flow controllers. The composition of the air was then checked on the PrimaBT Mass Spectrometer (Thermo Scientific, Waltham, MA, USA) to ensure the right ratios were achieved. Five residence times passed without the sparging of enriched CO_2_ air. Then two residence times were taken between shifting 0 to 10, 20, 30 40, 50 and finally back to 0% CO_2_. Throughout the experiment the sparging flow was adjusted to ensure aerobic conditions at a DO of 25%.

### Analytical methods

The UltiMate 3000 HPLC (Thermo Scientific, Waltham, MA, USA) with 9 mM sulfuric acid as the mobile phase was used to quantify and detect the levels of organic acids and the carbon source. The quantification of compounds was performed based on the refractive index (RI) chromatograms measured on the Refractomax 521 detector (Thermo Scientific, Waltham, MA, USA).

The cell dry weight (CDW) was determined by drying known sample volumes after washing the cell pellet twice with a 0.9% NaCl solution. The samples were placed in pre-dried and weighed Eppendorf tubes and dried in an oven at 80 °C until a constant weight was reached.

For RNA sequencing, 1 mL of broth was sampled in pre-cooled Eppendorf tubes and centrifuged directly for one minute at 4000×g and 2 °C. The supernatant was discarded, and the remaining cell pellet was snap-frozen in liquid nitrogen. RNA extraction, quality control, library preparation, and paired-end sequencing were performed by an external service partner (BGI Europe, Copenhagen, Denmark).

### Transcriptome alignment, QC, and ICA

We searched the sequence read archive(Katz et al., 2022) for all samples annotated with the organism “*Yarrowia lipolytica* W29” and supplemented our own fastq files with all available data (65 samples).

Fastq reads for each sample were processed, aligned and quantified using the nf-core/rnaseq pipeline(Ewels et al., 2020). A three step QC was then performed on the assembled count matrix. First, samples that failed the following metrics based on the multiqc report were rejected: i) per base sequence quality, ii) per sequence quality scores, ii) per base n content, and adapter content. In the second step, hierarchical clustering was performed to identify samples that fell into outlier clusters, i.e., clusters with very few samples with unrelated conditions. In the third step, samples with a Pearson correlation of less than 0.9 between replicates were removed. The filtered count matrix was then log2 normalized and centered to a reference condition/sample to compute the log2 fold change of each gene from the reference condition. The normalized tpm was then used for running robust ICA and an optimal number of ICA components was determined(McConn et al., 2021; Sastry et al., 2019). Code for our quality control and ICA pipelines are available on GitHub: https://github.com/avsastry/modulome-workflow.

### BLAST against the *S. cerevisiae* TRN

To obtain a basic Transcriptional Regulatory Network (TRN) for enrichment of the independent components, a draft TRN was constructed for *Y. lipolytica* through a bi-directional blast on its protein sequences for each gene against *S. cerevisiae* for which some TRN information is available(Monteiro et al., 2020b; Teixeira et al., 2018). Only mappings with high similarity (E < 10^-4^) in sequence along with high coverage (>0.25) were considered. This presented a one-to-one mapping between *S. cerevisiae* genes and *Y. lipolytica* genes that results in a draft TRN for *Y. lipolytica* and seeing which sets of genes might be co-regulated by which transcription factor/regulatory element.

### iModulon characterization through text mining and ontology analysis

Each iModulons’ gene list was further analyzed to deepen the understanding of their biological relevance and overarching functionality. Lifelike^©^, a graph-based natural language processing tool, was used to identify relationships between genes using various public databases such as UniProt and KEGG(“Webpage,” n.d.). Then, geneontology.org, based on Panther DB, was used for ontology analysis. Results were combined to check for consistency and reach the best possible prediction of each iModulon’s functionality.

### Visualization with pymodulon

iModulonDB dashboards and visualizations were generated using the PyModulon package(Rychel et al., 2021; Sastry et al., 2021b).

## Results

### Generating iModulons for *Y. lipolytica*

Multiple relevant and potentially stressful large-scale fermentation conditions were tested in shaken and continuous bioreactor cultivations (Figure 1A). The RNA-Seq data obtained in this study was combined with all publicly available datasets to obtain the largest possible compendium of *Y. lipolitica* W29 transcriptomes. A total of 153 datasets passed quality controls (Methods). Note that nearly all publicly available data (55/65 samples) failed our quality checks, while only 17/160 samples from our lab failed.

**Figure 1.**
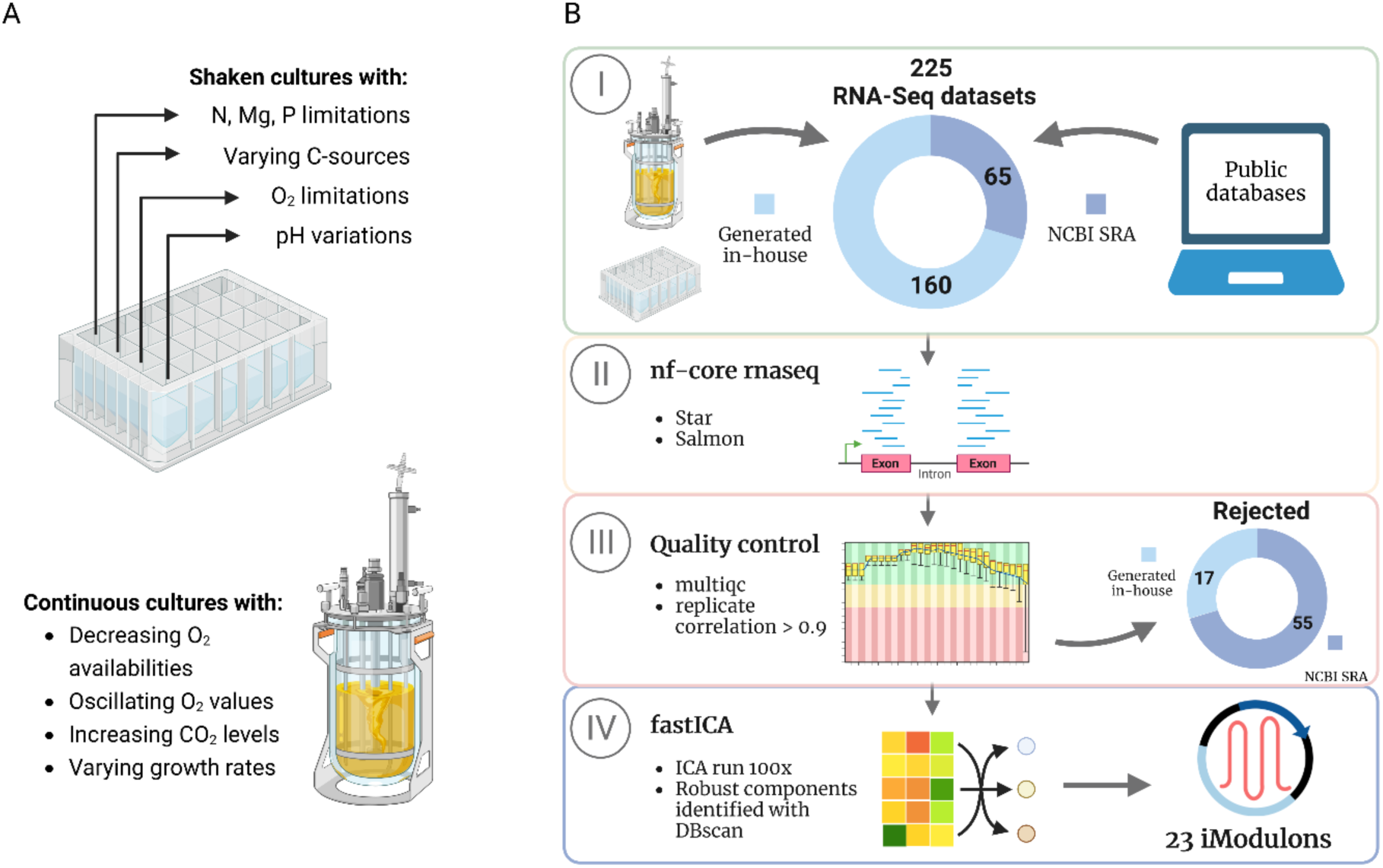
Schematic overview of the workflow adopted to identify independently modulated set of genes (iModulons) from the Y. lipolytica transcriptome. A. Overview of experimental conditions tested in this work. All systems were sampled to obtain RNA-Seq data and expand the compendium of transcriptomic datasets. B. Pipeline followed in this work to combine generated RNA-Seq data with publicly available datasets, alignment, quality control, and independent component analysis.

ICA of this data yielded 23 iModulons (Figure 1B). Each iModulon contains a set of genes that is independently modulated in all conditions present in the compendium. This list represents co-regulated genes. Often, a TF can be associated with an iModulon based on its known binding sites. Genes are either positively or negatively weighted in an iModulon, indicating an upregulation or downregulation of that gene. Then, in comparison to a reference condition, the iModulon activity can increase or decrease. This indicates how the activity of the associated TF and gene expression changes in response to environmental stimuli.

Given the lack of an existing knowledgebase for *Y. lipolytica*’s TRN, we mapped our iModulons to the genome of *S. cerevisiae* to aid in their characterization. We performed bidirectional BLAST between the two genomes, and then enriched the *S. cerevisiae* genes that correspond to each iModulon against the *S. cerevisiae* TRN available from YEASTRACT(Teixeira et al., 2018). BLAST showed good coverage of the *Y. lipolytica* gene set by *S. cerevisiae* genes (Figure 2A).

**Figure 2.**
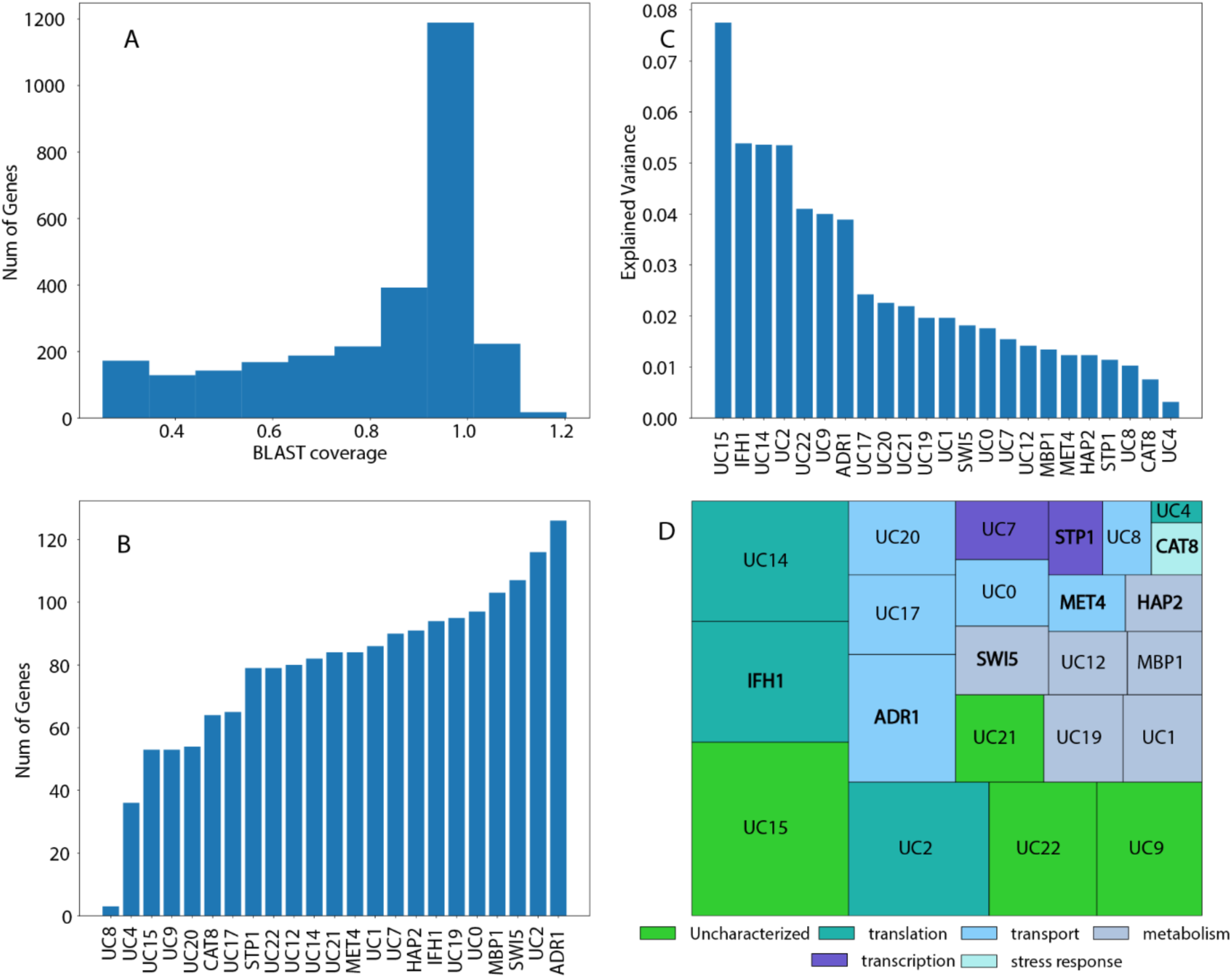
iModulon results and suspected functionality. A. Explained variance of the generated iModulons. B. Clustering of iModulon functionality based on ontology analysis and text mining of public databases. C. Blast coverage of conversion of Y. lipolytica into S. cerevisiae genes. D. Number of genes in each iModulon

Of a total of 23 iModulons, nine could be mapped against known *S. cerevisiae* regulons, and an associated transcription factor was identified. Thus, these iModulons were named based on their *S. cerevisiae* counterpart. For the remaining 14 iModulons, no statistically significant overlap with any *S. cerevisiae* regulon was found and thus they were named as ‘uncharacterized’ (UC). The iModulon with the largest explained variance in the compendium is Uncharacterized 15 (UC15) at 7.75% followed by IFH1 at 5.39%. Altogether, the 23 iModulons accounted for 60.2% of the explained variance (Figure 2B). The total number of genes represented by these the iModulons is 1262, which accounts for 19.5% of the total genome. The number of genes in an iModulon ranges from 3 to 126 genes with UC2, ADR1, MBP1, and SWI5 containing more than 100 genes (Figure 2C).

Text mining of various public databases and ontology analysis gave a suspected functionality and localization for most iModulons (Supplementary material). Confidence levels varied across the predicted functionalities, but they could be grouped into broad categories (Figure 2D). This analysis allowed for the clustering of six distinct functional groups of iModulons: six are associated with transport, four with translation, six are involved with metabolism, two with transcription, one with a general stress response, and four iModulons remain uncharacterized.

Interestingly, the literature description of some of the annotated iModulons does not align with the results found in the ontology analysis. The iModulon STP1, for example, has been associated with an extracellular amino acid sensing pathway, SPS, whereas ontology analysis linked it to microtubule motor activity (p-value 1.76E-07) and general mitosis(Omnus and Ljungdahl, 2014). While these discrepancies could be due to a variety of reasons, they are most likely determined by the following factors. First, in the conversion of *Y. lipolytica* genes to the *S. cerevisiae* regulon several conversion steps are made, which result in a loss of information. Second, due to the limited number of datasets, not all TRN dynamics are covered in the same detail. ICA identifies signals which appear to move as a unit in the transcriptome. As a result, different regulons which are co-stimulated under all conditions can be grouped into one iModulon that contains a high number of genes. As more diverse conditions are added to the compendium, the response of the TRN is further elucidated, causing iModulons to split into two or more new groups(Sastry et al., 2021a). Lastly, one of the complex factors of applying ICA to eukaryotic strains is the presence of various impactful global regulators. Their interaction with genes is often complex as they can vary between repression and activation, and are often not singular and linear in their activity. Considering all these factors, the functionality obtained from ontology is used for further interpretation in this work. Despite these limitations, the iModulon structure represents the most complete global TRN for *Y. lipolytica* and reveals important insights into regulatory functioning in this industrial biomanufacturing host.

Information on each iModulon, including its member genes, activity levels across all conditions, and the concordance between it and its matched *S. cerevisiae* regulon is available to browse and search on iModulonDB.org (Rychel et al., 2021).

### Clustering reveals correlated iModulon activities related to growth rates

iModulon activities were clustered based on the correlation between their activities across the entire dataset. Subcluster 11 (Ribosomes, metabolism, and transport) showed the best correlation across three different iModulons (Figure 3A). Two out of the three correlating iModulons were found to be uncharacterized but gene ontology analysis and text mining provided insights into their biological function. Interestingly, one of the iModulons is ADR1, which has a negative correlation with UC2 and a positive one with UC0. Consequently, UC2 and UC0 have a negative correlation as well.

**Figure 3.**
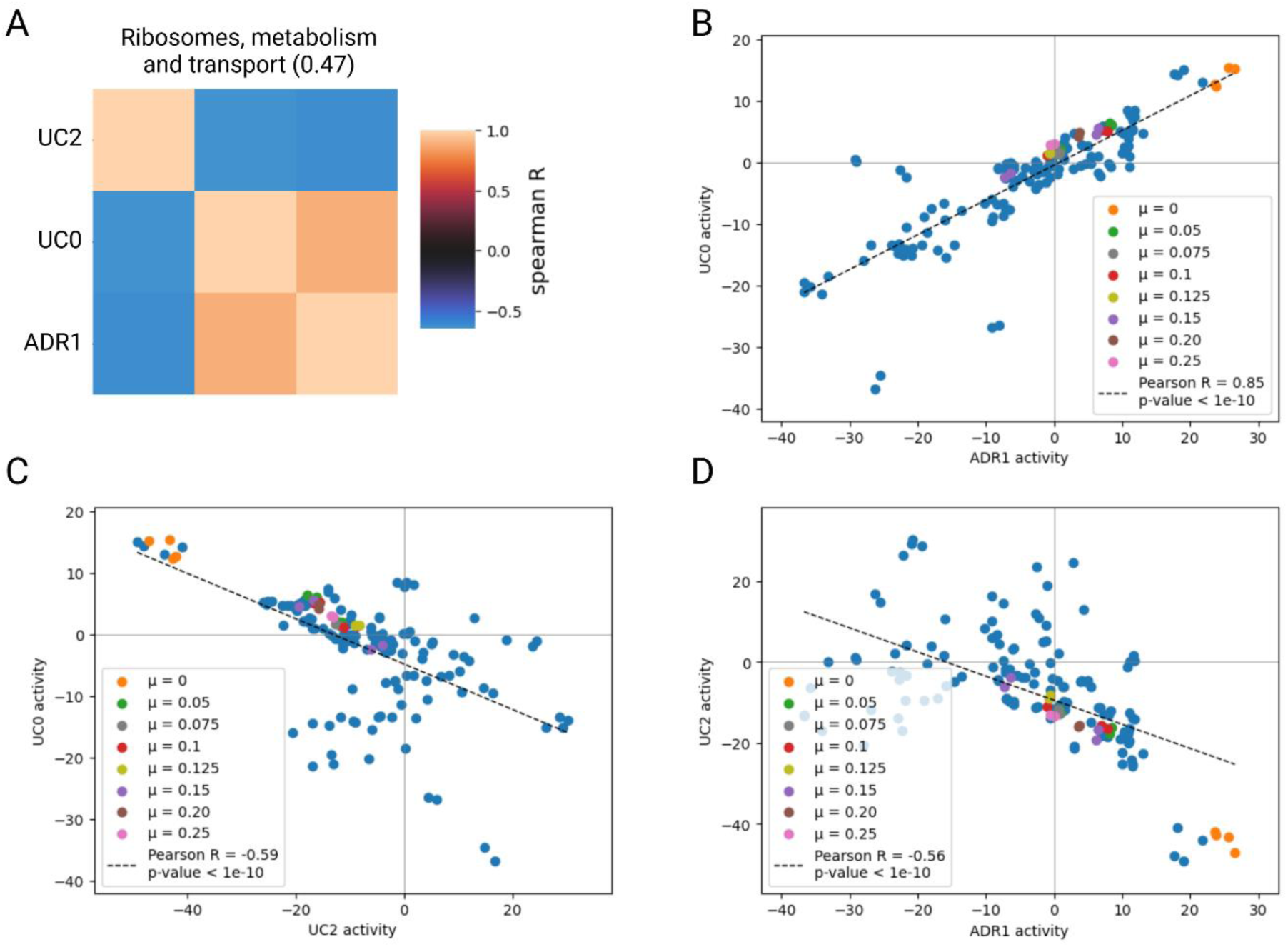
Clustering of iModulon activities. A. The cluster with highest correlation between iModulon activities across conditions contains three iModulons. B-D. Correlation between UC0 and ADR1, UC0 and UC2, and UC2 and ADR1. UC0 shows involvement with general transport, UC2 with ribosome biogenesis and ADR1 with general metabolism.

UC0 consists of 97 genes and appears to be involved in general transport with amino acid transmembrane transport (p-value 5.26E-05), organic acid transport (p-value 6.22E-06), and carboxylic acid transmembrane transport (p-value 1.05E-06) as primary functionalities. It has a positive correlation with ADR1 (Figure 3B). Biologically, this would be an understandable relationship as ADR1 activates the transcription of genes required for metabolizing non-glucose carbon sources. UC0 appears to be transporting non-glucose carbon sources into the cell, which could lead to a rearrangement of the cells’ machinery to adequately metabolize these compounds. The highest activity of both iModulons is found at the lowest growth rates. In this situation, where cells are deprived of glucose, it would be logical that cells scavenge their environment for alternative carbon sources.

UC2 contains 116 genes and shows a strong involvement in ribonucleoprotein complex biogenesis (p-value 1.08E-45) and ribosome biogenesis, specifically (p-value 1.79E-04). An increased activity of this iModulon correlates to a decreased UC0 activity. A higher UC2 activity would indicate an increased translational capacity, possibly linked to faster growth. Indeed, along the correlation trend line, UC2 activity is lowest for the non-growing cells and steadily increases as the growth rate increases (Figure 3C). A low ribosomal biosynthesis through UC2 is correlated to an increasing ADR1 activity (Figure 3D). It is possible that as the cells are metabolizing alternative carbon sources there is not sufficient energy being generated for further growth, resulting in the observed low UC2 activity.

### Elevated CO2 levels alter ribosomal synthesis and cell cycle progression

A continuous culture of *Y. lipolytica* was exposed to increasing levels of CO_2_ in the sparging air. As this increases the partial pressure of CO_2_ in the gas phase, a higher dissolved CO_2_ concentration will be reached. As a primary physiological performance indicator, the biomass yields, and glucose uptake rates were determined (Figure 4A). Both parameters indicate the same trend with an unaltered behavior up to a CO_2_ concentration of 20%, after which the yield decreases and specific uptake rate increases. As no significant by-products are measured under these conditions, it can be hypothesized that the elevated CO_2_ levels increase the maintenance requirements of cells. This is in line with previous research on *S. cerevisae* that saw a reduced biomass yield after prolonged exposure to elevated CO_2_ concentrations(Eigenstetter and Takors, 2017; Richard et al., 2014). As dissolved CO_2_ enters the cells, various intracellular changes occur, such as a decreased pH and change in ion balance(Buchholz et al., 2014). To counter these changes, cells need to increase their maintenance, causing a peak ATP demand and loss in energy. In systems with low pH values, these stresses are hypothesized to be relayed through the common (MAPK) signaling pathways(Hakkaart et al., 2020). As the elevated CO_2_ concentration is relieved, yields and uptake rates resume their original values, indicating a flexibility of the metabolic network(Kitano, 2004).

**Figure 4.**
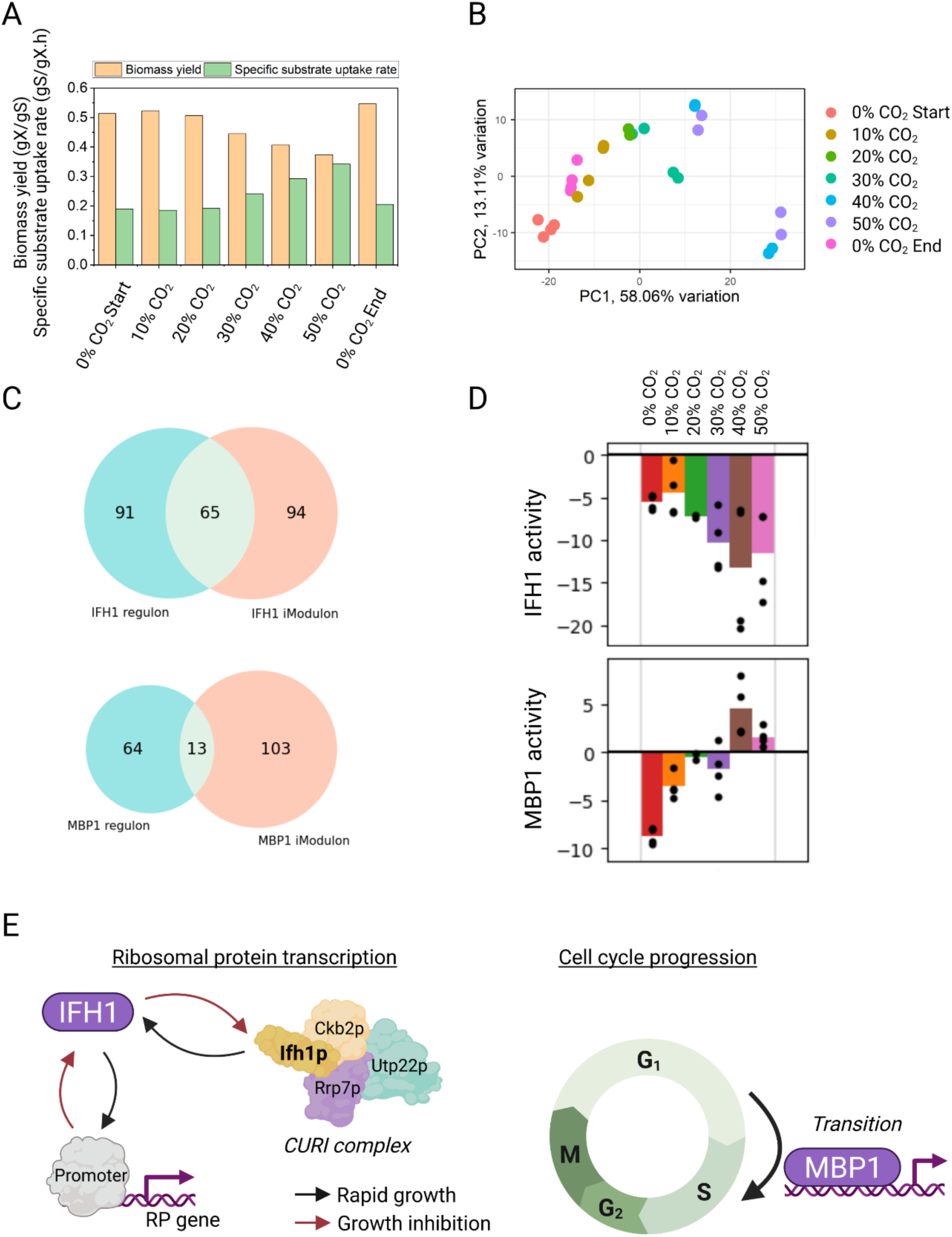
Transcriptional and regulatory changes in response to elevated CO_2_ concentrations. A. Biomass yields and specific substrate uptake rates across different steady state CO_2_ levels. B. Principal Component Analysis highlighting the general effect of elevated CO_2_ levels on the transcriptome. C. Venn diagrams of the IFH1 and MBP1 iModulon vs their corresponding S. cerevisiae regulons. D. Activities of IFH1 and MBP1 during exposure to increasing CO_2_ levels. E. Visualization of the biological functionality of the IFH1(Albert et al., 2016) and MBP1 transcription factors.

To further elucidate the response of a strain to varying (stressful) conditions, Principal Component Analysis (PCA) was performed on the transcriptomic data, yielding interesting profiles (Figure 4B). As CO_2_ levels increase, data points move across PC1, and nearly comparable values are obtained when returning to a CO_2_ percentage of 0. However, what is often found in these efforts are the limited interpretation possibilities. The two principal components with the highest explained variance provide information on the general expression of genes given certain conditions. The PCA analysis could be followed up with a volcano plot and gene ontology analysis, but this does not provide a systems level understanding. Thus, besides revealing (highly) up- or downregulated pathways, the underlying mechanisms remain hidden.Since iModulons are much more interpretable and more likely to match the biological transcriptional regulatory mechanisms, iModulon analysis has the potential to provide significant additional insights. In relation to the CO_2_ stress, two iModulons appeared to be of special interest.

The IFH1 iModulon had the best recall and precision values in comparison to its regulon (Figure 4C). As the CO_2_ concentration increased, the activity of this iModulon decreased (Figure 4D). This observation is biologically relevant as the transcription factor IFH1 is a coactivator that regulates the transcription of ribosomal proteins (RP) (Figure 4E). Through its product, Ifh1p, IFH1 also has an interesting interaction with the CURI complex(Albert et al., 2016). Under long-term stressful conditions, this complex would inhibit IFH1 activity, thereby orchestrating pre-RNA processing and RP production. As CO_2_ levels increase, with decreasing biomass yields as a result, it can be expected that ribosomal biosynthesis is decreased accordingly. Eukaryotic RP genes are actively regulated in response to various environmental stresses, which allows them to adjust total ribosome numbers and translation capacity accordingly(Wade et al., 2004). As ribosomes are crucial for protein synthesis in all known organisms, rRNA maintenance and transcription are tightly regulated to satisfy the required copy number(Hori et al., 2023). Moreover, ribosome production accounts for up to 90% of transcription in growing cells(Martin et al., 2006). Thus, the decreased yield observed (Figure 4A) can be partially explained by a decrease in RP gene expression (Figure 4D) and linked to IFH1 regulation through an analysis of this iModulon.

The MBP1 iModulon also displays interesting activity profiles (Figure 4D). With increasing CO_2_ levels, its activity increases and peaks at a CO_2_ level of 40%. After that, at CO_2_ 50%, the MBP1 activity decreases again. MBP1 has been found to be involved in the regulation of cell cycle progression from the G1 to S phase(McIntosh et al., 2000). The G1/S transition is primarily regulated by three main transcription factors, SWI4, SWI6, and MBP1. Together they form two complexes (SBF and MBF) that act through clearly distinct mechanisms on largely non-overlapping sets of genes(Harris et al., 2013). As the increasing CO_2_ level leads to an increased fraction of the cell population experiencing stressful conditions, the cell cycle can be disturbed. The G1 phase is a crucial part of the cell cycle and focuses on cell growth and the multiplication of organelles such as ribosomes. The cell cycle is a complex mechanism and a shortened G1 phase has been linked to an alteration in cell size(Talia et al., 2007; Turner et al., 2012). In filamentous fungi, a cell cycle arrest has been linked to filamentous growth(Chen et al., 2018). As *Y. lipolytica* has similar dimorphic properties, the MBP1 iModulon could be an indicator of filamentous growth. As such, the progression of the G1 phase is an important indicator of various cellular processes. Interestingly, the drop in MBP1 activity at a CO_2_ level of 50% coincides with a glucose accumulation in the reactor. This accumulation indicates that stresses have become so severe that the culture is no longer able to fully metabolize all carbon provided. Consequently, cells transition into the stationary or perhaps even decay phase so MBP1 becomes less active. Thus, the MBP1 iModulon acts as an indicator for the G1 phase of growth, and matches expectations based on growth rate and glucose uptake across this time course.

### O2 availability alters respiratory activity and redox homeostasis

In a repeated oxygen perturbation experiment, samples were taken frequently to assess the changes of the transcriptomic state. After the initial perturbation, the dissolved oxygen concentrations were continuously oscillated through an aerobic and anaerobic phase for a period of 10 hours. During the last oscillation, the bioreactor was again sampled during an aerobic and anaerobic phase (Figure 5A). The iModulon activity was monitored over the experiment duration and similar to the elevated CO_2_ experiment, IFH1 shows a decreased activity while MBP1 increases. This effect appears to be strongest during the initial perturbation with lower impacts after repeated oscillations. The strongest response, however, is found in the decreased activity of the ADR1 iModulon. This transcription factor is found to be involved in central carbon metabolism, especially with genes that are regulated by glucose repression. In the absence of glucose, Adr1p activates the transcription of genes required for metabolizing non-glucose carbon sources, such as non-fermentable carbon sources and fatty acids. An elevated ADR1 activity is measured for the first sampling point after which it decreases during the oxygen perturbations. The first sample is taken during a C-limited chemostat and as the oxygen oscillations start, repeated glucose accumulation and consumption is observed. Consequently, there could be a reduced trigger to deploy the ADR1 iModulon, which corresponds to the activity profile observed. Regulation by ADR1 is obstructed through phosphorylation by cyclic AMP-dependent protein kinase(Cherry et al., 1989). Anaerobic conditions interrupt oxidative phosphorylation with a decrease in ATP and increase in AMP and ADP as a result. Consequently, cyclic AMP-dependent protein kinase could be activated, resulting in a decreased ADR1 activity. After repeated oscillation, however, this effect is reduced considerably, indicating a rearrangement in the cells’ signaling and regulatory network. Thus, the desensitization of TRN activity by repeated oscillations is observed and quantified by iModulons, and particularly affects the ADR1 regulon. Other iModulon activities in the time course are presented in Figure 5A. Aside from this observation, HAP2 shows a downregulation only after repeated oscillations but none of the other characterized iModulons appear to be severely affected.

**Figure 5.**
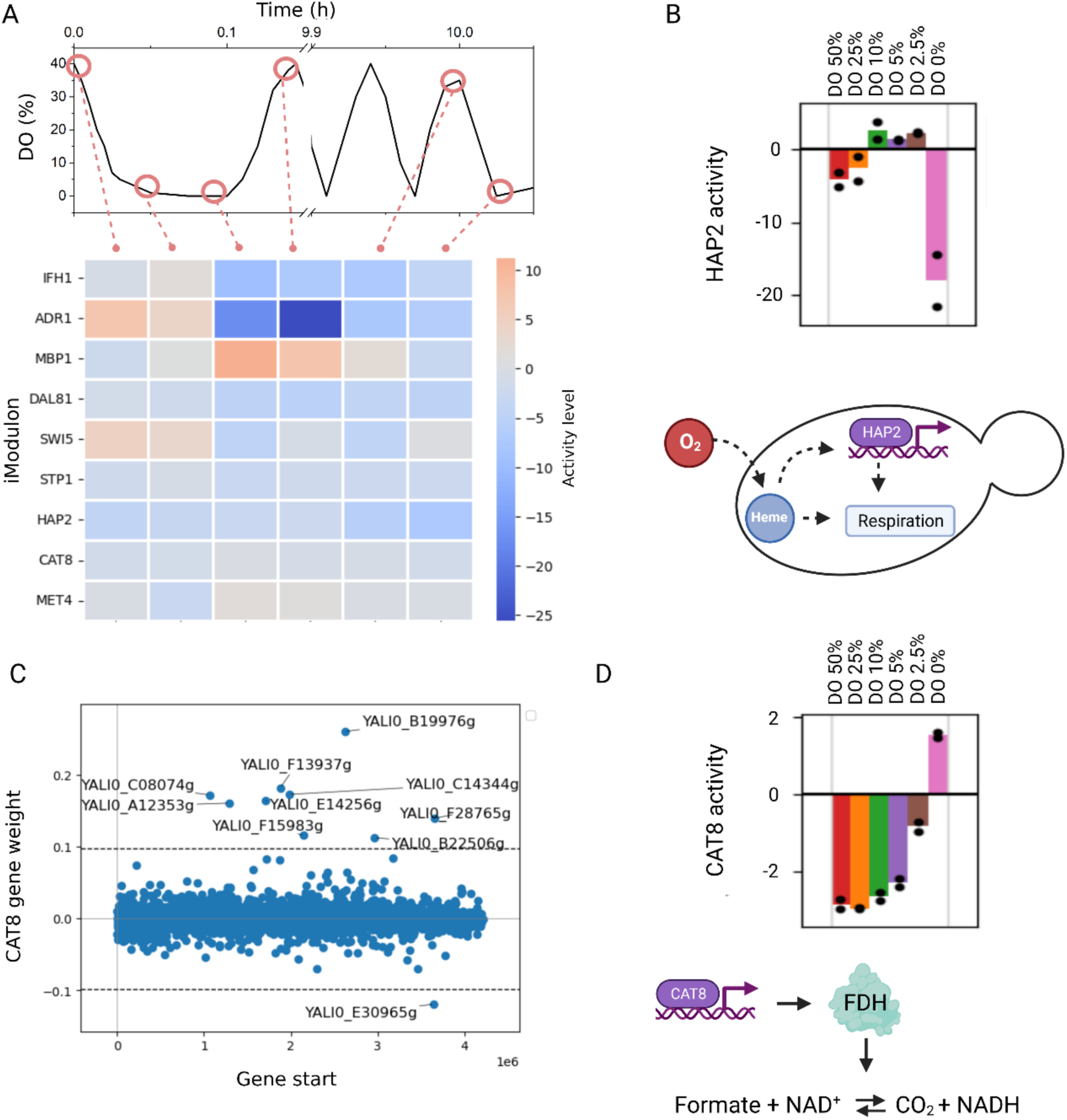
iModulon activity in response to varying oxygen availabilities. A. iModulon activity during an industrially relevant oxygen oscillation profile. Over the course of the first oscillation, the bioreactor is sampled frequently. When aerobic conditions are restored, a continuously oscillating profile is started. Cells cycle through aerobic and anaerobic conditions in a period of 5 minutes for a total of 10 hours. Then the bioreactor is again sampled during an aerobic and anaerobic phase. B. Activity and biological interaction of iModulon HAP2. C. Weights of all Y. lipolytica genes in relation to the CAT8 iModulon. All genes above and below the cutoff values (dotted lines) are considered to be significantly contributing and are thus part of the iModulon. All positively contributing genes are formate dehydrogenases. D. Activity and biological interaction of iModulon CAT8.

The compendium also contains a group of experiments assessing the effects of various steady state DO values ranging from 50% to approximately 0%(Kerssemakers et al., 2023). Both HAP2 (80 genes) and CAT8 (53 genes) showed strong responses as the dissolved oxygen concentration decreased, with an especially clear activity change for the system at a DO of 0% (Figure 5B). HAP2 is a subunit of the HAP2/3/5 heme-activated and glucose repressed complex. It is a transcriptional activator and global regulator of respiratory gene expression binding to the CCAAT sequence located on almost all cytochrome genes(Mao and Chen, 2019). Along with heme, HAP2 has been shown to be closely involved in the expression of COX6, subunit VI of cytochrome c oxidase. Heme synthesis correlates to the oxygen tension in the environment which triggers the HAP complex which in turn activates the transcription of genes involved in respiration(Kwast et al., 1998; Zhang and Hach, 1999). This activation happens through a delicate network and is mediated through the inactivation of ROX1 by heme, which encodes a repressor of many hypoxic genes(Zhang et al., 2017).

Interestingly, for CAT8, all significant positively weighted genes (YALI0B19976g, YALI0C14344g, YALI0F13937g, YALI0E14256g, YALI0C08074g, YALI0A12353g, YALI0F28765g, YALI0F15983g, YALI0B22506g) encode formate dehydrogenases (FDH) (Figure 5C). FDH has been shown to play a crucial role in anaerobic respiration in yeast, as it aids in redox homeostasis (Figure 5D). FDHs are responsible for catalyzing the reversible reaction between formate and CO_2_ (formate + NAD^+^ ⇌ CO_2_ + NADH + H^+^). As the TCA cycle is obstructed during anaerobic conditions, alternative NADH generating pathways could become more relevant. Negatively loaded on this iModulon is Acetyl-CoA hydrolase (ACH1), linked to the conversion of acetyl-CoA to acetate. Interestingly, upregulation of FDH in *Y. lipolytica* was also found to correlate with lipid accumulation and fatty alcohol production(Dahlin et al., 2019; Zhang et al., 2019). Additionally, formate has been shown to be involved with energy metabolism by promoting the synthesis of adenine nucleotides(Oizel et al., 2020). As energy availability drops due to decreased ATP concentrations under anaerobic conditions, formate appears to also act as an important signaling molecule. Interestingly, for the elevated CO_2_ experiments, CAT8 activity decreases with higher CO_2_ levels. This could thermodynamically favor the reverse FDH activity leading to the production of formate. To prevent formate accumulation, cells might upregulate this iModulon’s activity. Thus, varying oxygen availabilities appear to trigger signaling pathwaysinvolved in both respiratory pathways and energy homeostasis through iModulons HAP2 and CAT8.

## Discussion

In this study, we presented the first top-down omics data-driven elucidation of a quantitative TRN for *Y. lipolytica*. The study represents the first use case of iModulon analysis for a eukaryotic organism. We characterized iModulons by homology-mapping to the known TRN of *S. cerevisiae*, revealing the effects of nine characterized yeast transcription factors on this dataset. We propose a knockout study on the homologs of these transcription factors in *Y. lipolytica* to confirm the suspected regulation. Moreover, we reveal a cluster of iModulons, which exhibit correlated changes and provide context for uncharacterized iModulons. We then demonstrate the usefulness of iModulons for summarizing transcriptional dynamics under industrially relevant conditions. We find that the IFH1 and MBP1 iModulons cause a downregulation of ribosomal protein biosynthesis and activation of cell cycle progression genes under elevated CO_2_ concentrations. The ADR1 iModulon facilitates changes in carbon source uptake during a DO perturbation, but those changes are less extreme after repeated oscillations. Moreover, we found that the HAP2 and CAT8 iModulons are involved in respiration and energy homeostasis regulation under reduced oxygen availabilities. Our results demonstrate that iModulon analysis can be applied to eukaryotic transcriptomes and poorly characterized organisms, and it reveals the function of major transcriptional regulators that control phenotypic expression of *Y. lipolytica*.

ICA has proven to be very useful to study the TRN for a broad range of prokaryotic organisms(Rychel et al., 2021). Prokaryotic genomes are typically smaller in size compared to eukaryotes and are, in many instances, also better annotated. Consequently, their TRNs are often better known, which allows for a detailed interpretation of ICA results. This research is the first of its kind for eukaryotic strains. Due to different molecular mechanisms such as alternative splicing, the lack of operons, and heavy phosphorylation, regulation of gene expression is more complex for eukaryotes. Additionally, transcriptional regulation is often not singular or linear with a variety of global regulators interactively regulating the expression of the same gene(Balaji et al., 2006). Despite these complications, ICA revealed meaningful signals in the *Yarrowia* dataset. Therefore, we encourage continued data generation and development of advanced matrix decomposition algorithms to advance the development of novel mathematical frameworks for the analysis of eukaryotic transcriptional regulation.

The 23 iModulons identified here explain 60.2% of the total explained variance and, therefore, do not capture the full complexity of *Y. lipolytica* transcriptional regulation. Similar efforts on prokaryotic strains resulted in a higher number of iModulons and better explained variance. For example, for *Pseudomonas putida,* 84 iModulons explained 75.7% of the variance, for *Pseudomonas aeruginosa* 104 iModulons explained 66% explained variance, and for *Bacillus subtilis* 83 iModulons explained 72% explained variance, and all yielded more detailed results for the respective TRNs(Lim et al., 2022; Rajput et al., 2022; Rychel et al., 2020).

In part, these differences can be explained by the size of the transcriptome compendia used. For example, in PRECISE 1K, the latest transcriptomic compendium for *E. coli*, a total of about 1000 samples was used(Lamoureux et al., 2022) whereas this work is limited to 225 samples. Along this increased number of samples comes a variation in environmental conditions tested. Different conditions will result in an altered cellular behavior, thereby triggering different parts of the TRN. This enables ICA to capture more independent signals in the compendia, thus increasing the number of iModulons and potentially decreasing the number of genes in each iModulon. The results in this study show that there are several large iModulons that are often also larger in size than their *S. cerevisiae* regulon counterparts. It is likely that these iModulons capture genes from multiple regulators, whose individual TFs were not separately differentially activated by any conditions in the dataset. The addition of new samples could lead to one iModulon splitting into two or more, thereby proving a more accurate depiction of the *Y. lipolytica* TRN.

Another challenge is the lack of annotation of the *Y. lipolytica* iModulons. Therefore, to link the expression profiles to any known regulators, parallels had to be drawn to the *S. cerevisiae* regulon. This introduces a couple of limitations to this work. First, *Y. lipolytica* genes had to be blasted against *S. cerevisiae* to obtain the required annotation. Due to the differences between the two genomes and technical limitations related to the BLAST approach, this conversion results in the loss of relevant genes. In this work, this meant a loss of approx. 57% of the genes in our enrichment analysis. Secondly, the *S. cerevisiae* regulons are not fully annotated and validated with a total coverage of approx. 35% of the total genome. Although these factors impact the TRN annotation, they do not affect the outcome of ICA. To further build on these results, the *Y. lipolytica* regulons should be better annotated. Often, work on the TRN in *Y. lipolytica* focusses on the effect of a single TF through knockout experiments or the response to a single environmental stressor(Hirakawa et al., 2009; Martinez-Vazquez et al., 2013; Pomraning et al., 2016). Other efforts might deploy a larger TF screening efforts and focus on a single performance parameter such a lipid accumulation(Leplat et al., 2018; Trébulle et al., 2017). These contributions are of great value to feed the ICA pipeline and validate the iModulon results.

Taken together, this initial study applying ICA to eukaryotic transcriptomes was successful. We identified physiologically meaningful iModulons that represented key cellular processes and thus phenotypes. iModulon activities under bioprocess conditions can thus represent a useful approach to process optimization. Our results provide a strong impetus for undertaking more comprehensive studies of iModularization of eukaryotic transcriptomes and determining their utility under real-life bioprocessing conditions.

## Acknowledgements

We thank the yeast metabolic engineering group at the Novo Nordisk Foundation Center for Biosustainability for supplying the strains used in this research. Some of the visuals in this work were partially created through Biorender.

## Data availability

RNA-Seq files have been uploaded to SRA under BioProject numbers PRJNA949438, PRJNA949440, and PRJNA949441. iModulon dashboards and downloadable processed data are available at iModulonDB.org under the organism “*Y. lipolytica*” and dataset “YarrowiaPRECISE”.

## Author contributions

**Abraham A.J. Kerssemakers:** Conceptualization, Methodology, Software, Validation, Formal analysis, Investigation, Data Curation, Writing - Original Draft, Writing - Review & Editing, Visualization, Funding acquisition. **Jayanth Krishnan;** Conceptualization, Methodology, Software, Validation, Formal analysis, Investigation, Data Curation, Writing - Original Draft, Writing - Review & Editing, Visualization. **Kevin Rychel**: Conceptualization, Methodology, Software, Formal analysis, Data Curation, Writing - Review & Editing. **Daniel C. Zielinski**: Conceptualization, Methodology, Software, Writing - Review & Editing, Supervision, Project administration. **Bernhard O. Palsson**: Conceptualization, Methodology, Writing - Review & Editing, Supervision, Project administration. **Suresh Sudarsan:** Conceptualization, Methodology, Writing - Review & Editing, Supervision, Project administration, Funding acquisition.

